# Selenium in wheat from farming to food

**DOI:** 10.1101/2021.07.17.452805

**Authors:** Min Wang, Baoqiang Li, Shuang Li, Ziwei Song, Fanmei Kong, Xiaocun Zhang

## Abstract

Selenium (Se) plays an important role in human health. Approximately 80% of the world’s population does not consume enough Se which recommended by WHO (World Health Organization). Wheat is an important staple food and Se source for most people in the world. This article summarizes literatures about Se from 1936 to 2020 to investigate Se in wheat farming soil, wheat, and its derived foods. Se fortification and the recommended Se level in wheat were also discussed. Results showed that Se contents in wheat farming soil, grain, and its derived foods around the world were 3.8–552 (mean, 220.99), 0–8,270 (mean, 347.30), and 15–2,372 (mean, 211.86) μg·kg^−1^, respectively. Adopting suitable agronomic measures could effectively realize Se fortification in wheat. The contents in grain, flour, and its derived foods could be improved from 93.94 to 1,181.92, 73.06 to 1,007.75, and 86.90 to 587.61 μg·kg^−1^ in average after leaf Se fertilizer application in the field. There was a significant positive correlation between Se content in farming soil and grain, and it was extremely the same between foliar Se fertilizer concentration rate and grain Se increased rate. The recommended Se fortification level in cultivation of wheat in China, India, and Spain was 18.53–23.96, 2.65–3.37, and 3.93–9.88 g·hm^−2^ respectively. Milling processing and food type could greatly affect Se content of wheat derived food and should be considered seriously to meet people’s Se requirement by wheat.

**TOC graphic:** 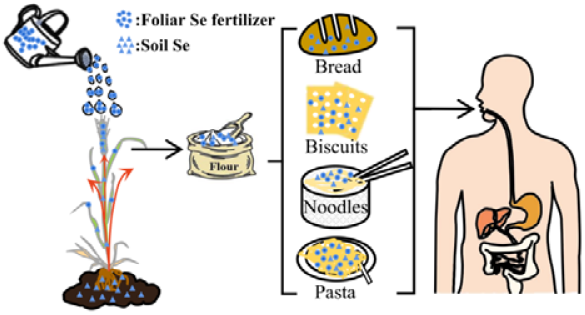

---

Selenium (Se) is a metalloid element of group VI A in the fourth period of the periodic table of the elements. It was first discovered by Swedish scientist Berzelius JJ in 1817 and named it after the Selene, the goddess moon^1^. However, researchers discovered in 1934 that the “alkaline soil disease” and “stumble” of animals were caused by excessive intake of Se. Since then, research on Se has mainly been based on its toxicity^2^. In 1860s, Se was determined as an essential element in animals and human bodies and has gradually gained widespread attention worldwide^3^. Se is one of the 14 essential trace elements in human body which constitutes a series of important Se-containing proteins and enzymes in galacturids^4^ although the total amount of it in the human body is only 14–20 mg^4^. Se is closely related to the health and Se deficiency can cause more than 40 kinds of diseases, such as Keshan disease, large bone nodule disease, anemia, infertility, and muscular dystrophy^6^. Moreover, excessive Se intake can cause hair and nail loss, skin ulceration, nerve damage, and other toxic symptoms^7^. Therefore, a safe and suitable daily Se intake should be maintained and 50–400 μg/day of Se intake is safe and suitable according to the recommendations of World Health Organization (WHO)^8^. Se deficiency is widely observed in more than 40 countries around the world, and approximately 80% of the population is Se deficient^9^. The most direct source to get Se is through dietary intake though Se bioavailability in human body varies widely (10%–85%)^10^. Therefore, improving the Se deficient situation of people through safe Se-rich diet is an effective and long-term task.

Wheat is the second largest food crop in population consumption^11^, and it provides nearly 50% daily calorie intake in most developing countries and more than 70% calorie intake in rural areas^12^. Wheat has strong Se enrichment capacity. It can produce high-security organic Se by absorbing Se in soil or foliage and store them in grains^12^. Se is mainly combined with protein in wheat grain and more evenly distributed throughout the whole grain than other minerals^14^. Therefore, unlike other micronutrients, most of the Se found in wheat grains can be retained in the flour, rather than being lost to the bran and aleurone layer during milling. Hence, it is an effective way to improve Se intake of people through Se enriched wheat.

Se fortification of wheat is a comprehensive and multi-objective process. Generally, the final effects of wheat Se fortification (Se content in the food) can be significantly affected mainly by three processes, including grain Se enriching in the field, grain grinding, and wheat-derived food processing. This article summarized reports related to the Se fortification of wheat, and discussed the main influencing factors, possible regulation measures and some relatively safe and suitable reference during the above three processes. The data of Se bioavailability was also discussed.

## 2 MATERIALS AND METHODS

### 2.1 Data collection

The papers relation to Se of wheat grain, flour and wheat derived foods published between 1936 and 2020 were used to set up the database by a literature search using Wiley, ACS, Science Direct, Springer Link and China National Knowledge Infrastructure (CNKI). Search keywords included different combinations, “wheat” AND “Selenium” or “flour” AND “Selenium” or “food” AND “Selenium”. Selection criteria included: a) Field experiment: report wheat harvest time, wheat type and fertilization type, including soil pH and Se content; b) wheat milling: report Se fertilizer type and Se content in different components after milling; and c) Food processing: report the source of wheat flour, food type, and Se content in food.

### 2.2 Data Classification

In selected reports, we summarized these data from soil to grain, milling, then to food under different conditions. A total of 66 research papers and 1066 paired observations were obtained (Figure 1); Grain Se content in different countries came from 41 references (Table S1); Se content of different wheat milling components were also obtained under different peeling and flour milling processing were got in 20 papers (Table S2); Se content of six kinds of different foods under different flour processing were obtained from 26 reports (Table S3). Total of 117 paired observations for the effects of soil Se on wheat grain from 21 articles, and 188 paired data for the effects of different Se fertilizer on wheat grain from 21 articles were extracted.

**Figure 1.**
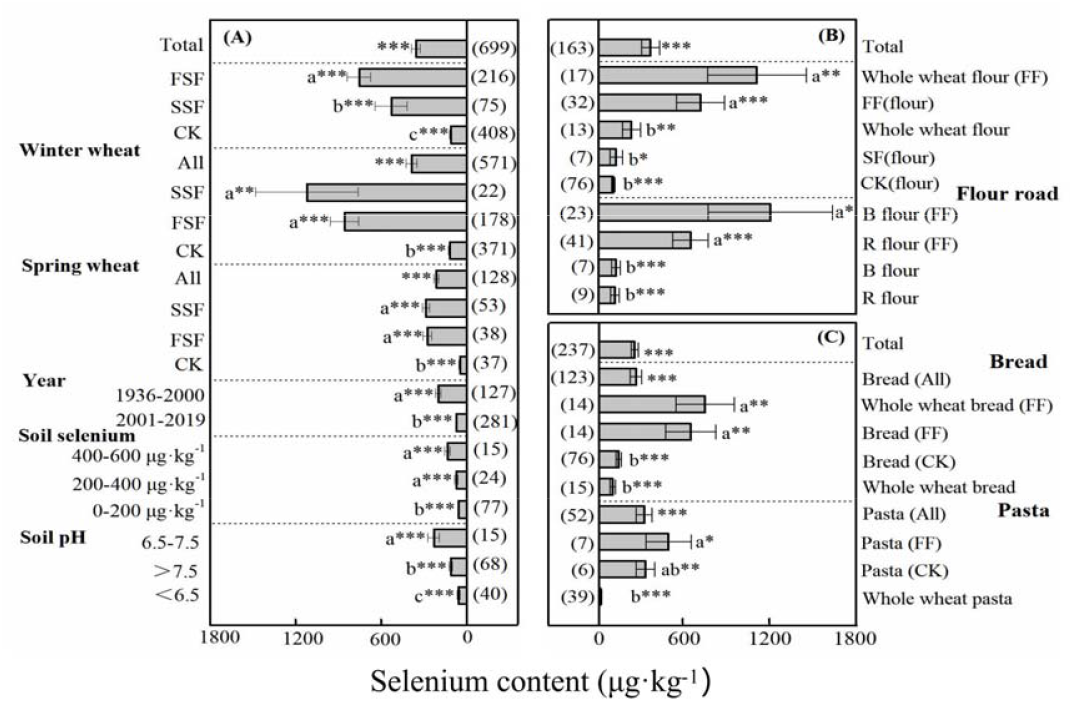
Comparison of Se content in wheat grain (A), flour (B) and food (C) **Note**: Data originated from 66 studies; FSF: foliar Se fertilizer treatment, SSF: Soil Se fertilizer treatment; CK: unfortified Se treatment; The year is based on the harvest year of wheat; Data are presented as mean ± 95% CIs, with n in parentheses. Asterisks indicate significant differences (\**p < 0.05, **p < 0.01, and ***p < 0.001*); Different letters indicate significant differences between mean effect size for groups within each category.

### 2.3 Data management

The database was established using Excel 2019 software. OriginPro 8.5 and PowerPoint 2019 software were used to draw graphics. All statistical analyses were performed using SPSS 18.0 software.

## 3 RESULT AND DISCUSSION

### 3.1 Se content in wheat grain

Se content in wheat grain was 0–1500 μg·kg^−1^ (110.98 μg·kg^−1^ in average) in the world^8, 15–50^. The average Se content was lower than that suggested by the reference of Se-rich grains (200–300 μg·kg^−1^) ^39^. The large variance of Se content in wheat grain was mainly attributed the difference of wheat varieties, area, basic properties of soil, and Se biological strengthening measures in the field.

#### 3.1.1 Variance of Se content among different wheat varieties

Generally, the old and new varieties might have different potentials to accumulate Se in grain. According to our data, grain Se content in wheat reported in 1936–2000 could reach 198.76 μg·kg^−1^ in average, and this value was significantly higher than that in 2001–2020 (Figure 1a). “Green Revolution”, which began in the 1950s, dramatically increased crop production^51^ and the “dilution effect” caused by the increase of yield might be the most important reason for the decrease of Se content in new varieties. In addition, different wheat varieties have different ability to accumulate Se under the same soil environment. For example, black grain wheat Se content was 177.34 μg·kg^−1^, which was significantly higher than that in white grain (70.29 μg·kg^−1^)^33^.

#### 3.1.2 Grain Se content of wheat in different continents and countries

No significant difference was observed among the average grain Se contents of wheat in Asia^8, 15–20, 22–25, 27–29, 31, 33–40, 43^ (63.36 μg·kg^−1^), Africa^8, 15–20, 22–25, 27–29, 31, 33–40, 43^ (64.15 μg·kg^−1^), and Europe^20, 22, 26, 29, 30, 41, 42, 44, 46, 48, 50^ (85.98 μg·kg^−1^), and these values which were all lower than the normal level (100 μg·kg^−139^). There was a significant difference among the grain Se content in South America^22, 50^, North America^20, 22, 47, 49, 50^, and Oceania^22, 45, 50^ with values of 600.00, 286.23, and 194.80 μg·kg^−1^, respectively (Figure 2). These values were significantly higher than that in Asia, Africa, and Europe. Grain Se content in different countries on the same continent varied greatly too. For Europe, the grain Se content in Spain, Slovakia, Portugal, and Norway were 26.75–93.90 μg·kg^−1^, but that in Italy was 133.44 μg·kg^−1^.

**Figure 2.**
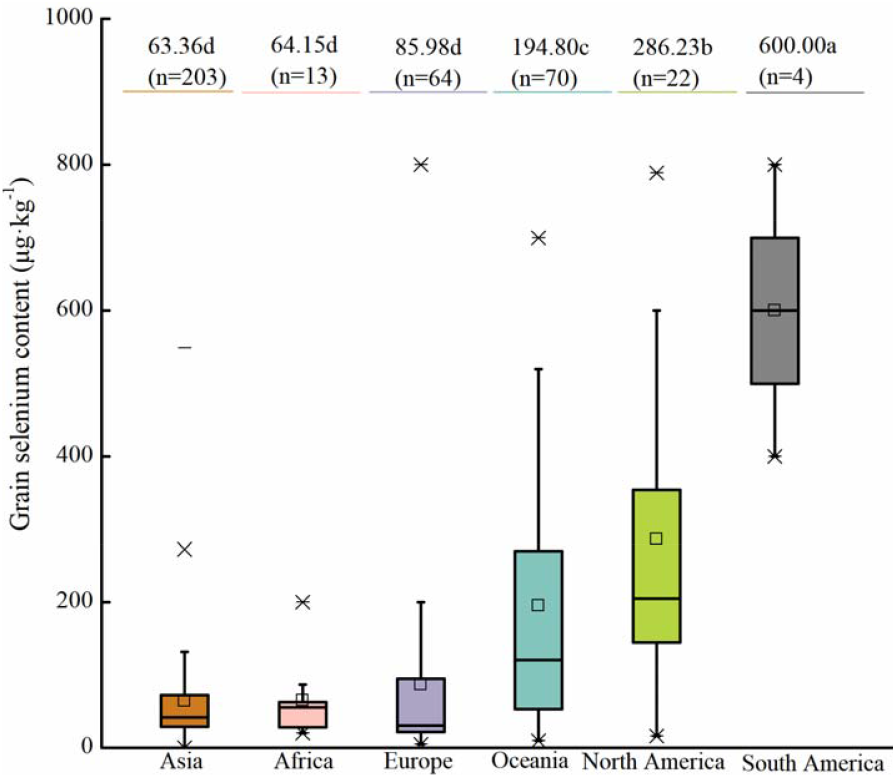
Selenium content of wheat among different continents **Note**: Data originated from 42 studies; The letters a, b, c and d indicate a significant difference at the level of P < 0.05; n is the total number of samples.

#### 3.1.3 Basic properties of soil affecting grain Se content of wheat

Soil Se content and pH are the basic properties of soil which can significantly affect Se content in wheat grain. The total soil Se content varied greatly among different regions. Se-deficient, Se-sufficient, Se-rich, and Se-high soil were considered at soil Se contents of 0–175, 175–450, 450–2,000, and 2,000–3,000 μg·kg^−1^, respectively^28^. The soil in most parts of the world is Se-deficient. For instance, soil Se contents in Italy, China, India, and South Africa are 130, 21.21 12.60, and 6.14 μg·kg^−1^, respectively. Globally, soil Se content in farming system is 10–2,000 μg·kg^−1^ (300 μg·kg^−1^ in average)^28^, while that in farming soil of wheat is 3.80–552.00 μg·kg^−1^. There was a significant positive correlation between soil Se content and grain Se content (Figure 3). Se content of wheat grain from the soil with Se content of 0–200 μg·kg^−1^ was significantly lower than that from the soil with Se content of 200–400 μg·kg^−1^ and 400–600 μg·kg^−1^, but no significant difference was observed between that from the soil with Se contents of 200–400 μg·kg^−1^ and 400–600 μg·kg^−1^ (Figure 1a).

**Figure 3.**
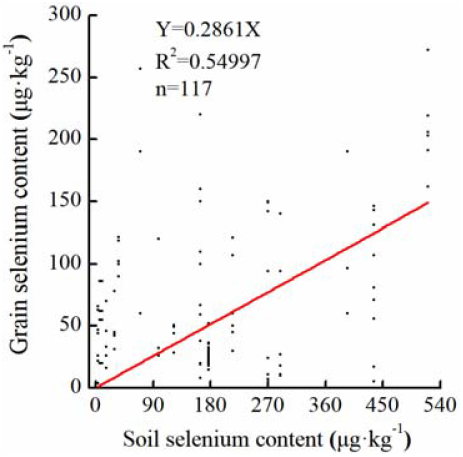
Relationship between soil selenium content and grain selenium content **Note**: Data originated from 21 studies; n is the total number of samples.

Soil pH also significantly affects grain Se content in wheat, and a significant difference was observed in wheat grain Se content among neutral soil (pH 6.5–7.5), alkaline soil (pH>7.5), and acid soil (pH<6.5). The grain Se content of wheat planted in neutral soil (pH 6.5–7.5) was the highest (229.42 μg·kg^−1^), whereas that in acid soil (pH<6.5) was the lowest (55.50 μg·kg^−1^) (Figure 1a). Acid soil occupies approximately 30% of the world’s ice-free land area and occurs mainly in tropical and subtropical regions. These acid soils are more likely to be Se-deficient^53^.

The Se content in soil is significantly positively correlated with the Se content in grain, so planting in Se sufficient soil is an important way to obtain Se-rich wheat. However, most cultivated soils in the world are Se insufficient, so applying Se fertilizer to enhance Se content in wheat grains is the preferred route to consider.

#### 3.1.4 Effects of different Se fertilization on grain Se content in wheat

The process of improving grain Se by fertilizer in the field is the most important part in Se fortification. Se fertilization treatment during wheat growth can significantly increase grain Se content according to the data about the grain Se contents in wheat around the world since 1936 (Table S1). The grain Se content of wheat planted in natural soil without Se fertilization treatment was about 0–1,500.00 μg·kg^−1^ (110.98 μg·kg^−1^ in average)^8, 15–50^, and that reached 58.00–8,270.00 μg·kg^−1^ (677.21 μg·kg^−1^ in average) with Se fertilization treatment.

However, significant differences were observed in the effect of different Se fertilizer treatments and most of the reports indicated that foliar Se fertilization treatment might be the most effective way to improve grain Se content^8, 15, 20, 26, 29, 35, 36, 52^. Grain Se content with foliar Se fertilization treatment was significantly higher than that of soil Se fertilization treatment and CK (Figure 1a). Se content in the wheat grain with soil Se fertilization treatment was 190.00–6,090.00 μg·kg^−1^ (528.87 μg·kg^−1^ in average)^19, 28, 29, 38, 42, 48, 54^, that with foliar Se fertilization treatment was 58.00–8,270.00 μg·kg^−1^ (729.69 μg·kg^−1^ in average)^8, 13, 15, 16, 18–20, 23–31, 35–38, 52, 54^, and that with seed Se soaking treatment was only 64.80–300.00 μg·kg^−1^ (128.67 μg·kg^−1^ in average)^39, 55^. A positive correlation was observed between foliar Se fertilizer application amount and grain Se content growth rate, and this correlation can be expressed with the equation *Y*=22.20587 *X* (Figure 4a). But there was no significant correlation between soil Se fertilization and grain Se content, which correlation can be expressed with the equation *Y=0.0157 X*. The use of foliar Se fertilizer was an environmentally safe and effective because of its low concentration in solution, effective absorption, and more accurate control of total Se application. Using 100 g Se·hm^−2^ for foliar fertilization, grain Se content is 1770 μg·kg^−1^; but the same Se dosage for soil fertilization, grain Se content is only 1064 μg·kg^−129^. The foliar Se fertilizer is the most popular method in Se fortification. Moreover, Na_2_SeO_3_ is the most common form of foliar Se fertilizer for its lower toxicity than Na_2_SeO_4_.

**Figure 4.**
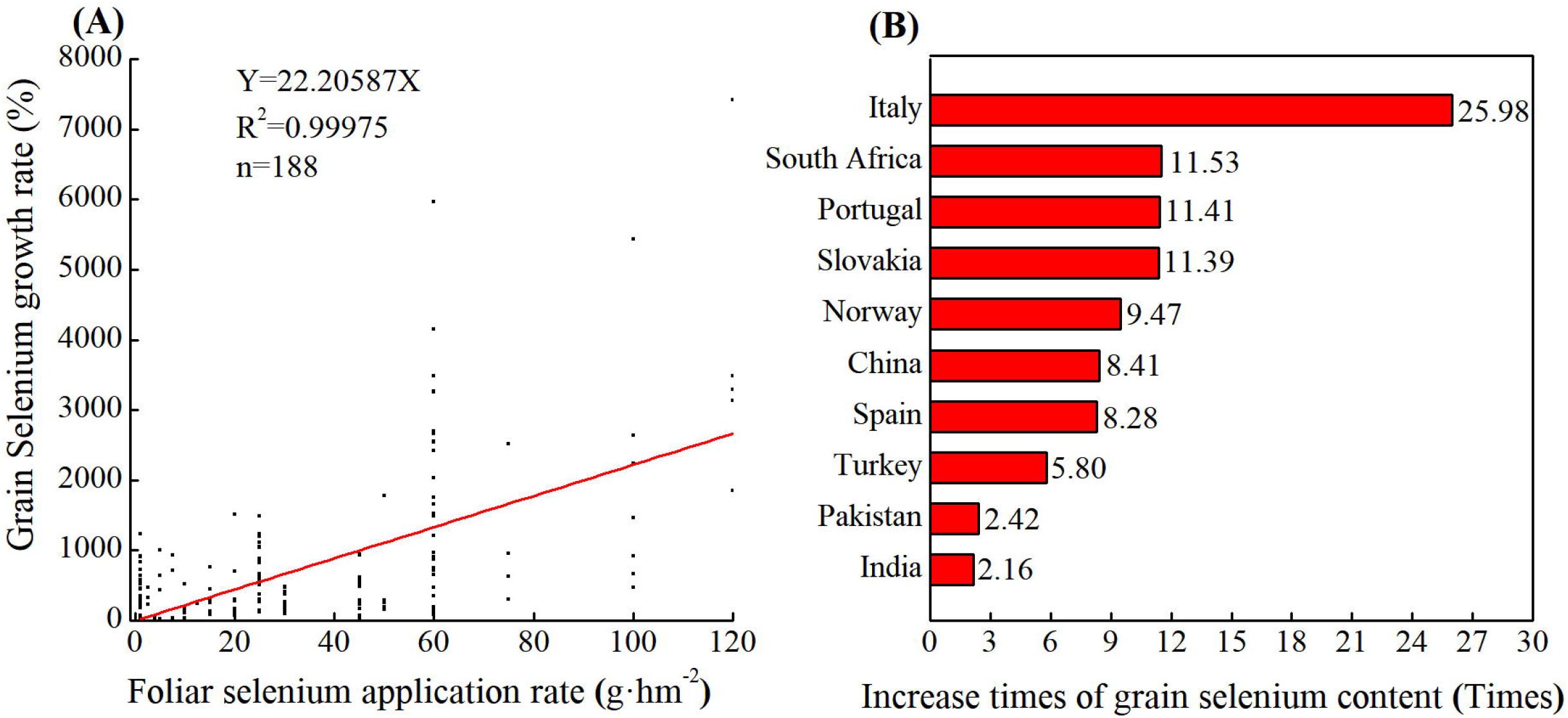
Relationship between foliar fertilization application rate and grain selenium content (A) and increase times of grain selenium content after fortification in different countries (B) **Note**: (A) Data originated from 21 studies; Selenium was added in the form of Na_2_SO_3_ solution; n is the total number of samples; (B) Data originated from 41 studies.

The grain Se content in different countries increased significantly after Se fortification in the field (Figure 4 b), and this value increased from 93.94 μg·kg^−1^ to 1181.92 μg·kg^−1^ after Se fortification. A large variance in Se fortification effects was observed in different countries. For example, the grain Se content increased in Pakistan, India, Turkey, Spain, South Africa, Norway, Portugal, Slovakia, China, and Italy by 2.16–25.98 times and that in Pakistan and Italy was 2.16 and 25.98 times, respectively.

In addition to the Se application amount, Se concentration is also an important factor affecting the effect of foliar fertilizer for one wheat variety. Wheat Se content can reach the standard of Se fortification by foliar fertilizer application (Se concentration: 40 mg Se·L^−1^, application rate: 750 kg·hm^−2^) at the early filling stage. Hence, the determination of the appropriate application concentration and amount of foliar fertilizer are both the key to the effect of Se nutrient enhancement of wheat.

### 3.2 Se in flour and its influencing factors

The proportion of Se in wheat grains that can be directly ingested by the human body is also a key link for human Se supplementation. Most of the wheat will be ground into flour, and then be made into different foods. Wheat grain is generally divided into bran, embryo, and endosperm, accounting for 14%–16%, 2%–3%, and 81%–84% of the grain weight, respectively^29^. During milling, wheat grains are peeled off. The bran, which consists of the seed coat, aleurone layer and embryo, are then stripped off. Se content in bran was 15.00–520.00 μg·kg^−1^ (149.48 μg·kg^−1^ in average)^13, 19, 26, 28, 45^, which was significantly higher than that in flour (101.02 μg·kg^−1^ in average)^13, 19, 26, 28, 30, 41, 45, 56–67^. There were also some reports indicated that Se content in bran is approximately three times higher than that in flour after grinding^19, 26, 45^. It was clear that, there should have a part of Se loss during the milling process.

Different wheat milling methods remarkably affect flour Se content (Figure 1 b). Flour can mainly be divided into two types, namely, whole wheat flour and standard flour, according to the milling method. Se content in standard flour was significantly lower than that in whole wheat flour. According to our results (Table S2), Se content in standard flour was 13.00–500.00 μg·kg^−1^ (101.02 μg·kg^−1^ in average)^13, 19, 26, 28, 30, 41, 45, 56–67^, and the whole wheat flour Se content was 35.60–650.00 μg·kg^−1^ (227.58 μg·kg^−1^ in average)^26, 57, 62, 64, 66, 68^. Flour road during flour milling can be divided into six flour parts (i.e., three “Broken [B]” and three “Reduced [R]” milling parts) and two kinds of bran parts (i.e., bran and shorts). A significant difference was observed among the ingredients in different flour roads. The flour collected by B1, R1, B2, R2, B3, and R3 was standard flour, the flour collected by B1, R1, B2, and B3 was bread flour, and the flour from B1 and R1 was refined flour^26, 57, 62, 64, 66, 68^. The Se content in different components after wheat milling can be arranged as follows: B flour >R flour. The Se contents of flour from R1, R2, R3, B1, B2, and B3 were 106.85, 128.32, 207.90, 89.00, 96.82, and 128.29 μg·kg^−1^ in average, respectively. The Se content of flour in common flour roads was B3>B2>R3>B1>R2>R1^57, 69^. The rate of Se concentration loss and Se content loss in wheat milling were 17.19% and 46.40%, respectively. Most of the published literature used Se concentration loss, and a few studies have focused on Se content loss. The peeling rate of 4%–5% can effectively increase standard flour Se content. When the peeling rate was greater than 9%, no significant difference was observed in the Se content between standard flour and unpeeled flour^19^.

Se content in standard flour and the products in each flour road can be greatly affected by grain Se content. Se contents in whole wheat flour and standard flour in the Se-rich grain with foliar Se fortification treatment were significantly higher than that in the control treatments, and increased from 227.58 μg·kg^−1^ to 1,110.03 μg·kg^−1^ and from 101.02 μg·kg^−1^ to 714.90 μg·kg^−1^ respectively (Figure 1b). The Se contents in the products of different flour roads for the Se-enhancing grain from foliar Se fortification could be arranged as B2>R3>B1>R2>R1, which was quite different with that of common flour. The average rates of Se concentration and content loss in wheat milling were 15.04% and 38.56%, respectively, after Se fortification treatment.

Therefore, milling methods might affect flour Se content, and foliar Se fortification can greatly improve Se content in standard flour and change the Se distribution in products of different flour roads.

### 3.3 Se in wheat derived foods

Flour is generally used for the preparation of foods, such as bread, noodles, and biscuits. These staple foods play an important role in people’s diet. For example, wheat-related traditional foods account for 75% of the total wheat consumption in China^34^. Se content in wheat-derived foods was 3.60–1130.00 μg·kg^−1^ with an average of 191.87 μg·kg^−1 13, 30, 41, 45, 46, 57, 59–62, 64–67, 70–81^ according to the data over the past 50 years (Table S3). There was a significant difference for the Se content of different wheat-derived foods (Figure 5). Cooking foods has the highest Se content which can reach 382.83 μg·kg^−1^, and it was significantly higher than that in baked foods (244.98 μg·kg^−1^). In addition, the average Se contents in pasta, noodles, biscuits, and bread also can achieve 325.45, 251.53, 181.65, and 137.51 μg·kg^−1^, respectively.

**Figure 5.**
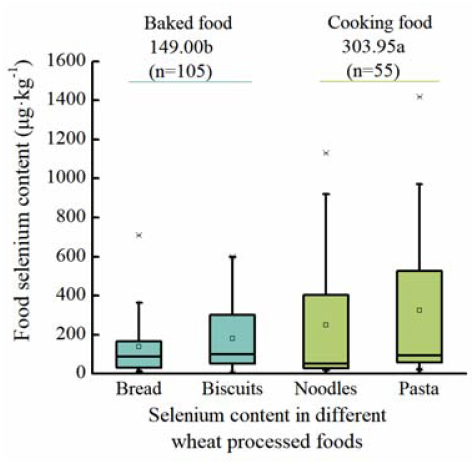
Selenium content in different wheat derived foods **Note**: Data originated from 26 studies; The letters a and b indicate a significant difference at the level of P < 0.05; n is the total number of samples.

**Figure 6.**
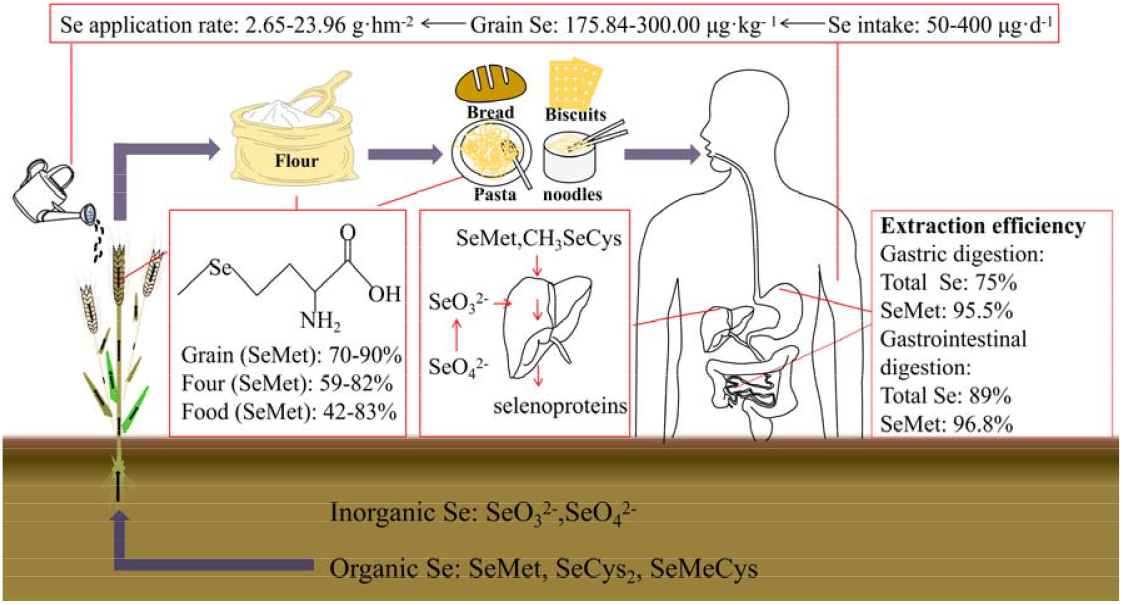
The change of selenium from grain to human body and the appropriate level of selenium fortification

High temperature can result in the volatilization of Se^83^. Hence, Se loss in wheat-derived foods processing is mainly occurs during heating, especially under the dual effect of dissolution and heat volatilization when cooking, steaming, and frying the same food^82, 83^. Moreover, soaking steps in food fermentation also can reduce wheat-derived foods Se content. For example, Se content in mung bean and millet would decrease by 7.17% and 2.62%, respectively during soaking^83^. But no report about the change of Se in wheat grain during soaking and the flour fermentation was retrieved.

In addition, increasing grain Se content through foliar Se fortification can effectively improve Se content in wheat-derived foods (Figure 1c). For example, Se content in pasta increased from 22.4 μg·kg^−1^ to 424.43 μg·kg^−1^ (17.94 times) and that in bread increased from 119.15 μg·kg^−1^ to 644.69 μg·kg^−1^ (4.41 times) using the flour derived from foliar Se fortification. Hart et al.^57^ found that foliar Se application rate is positively correlated with bread Se content, and their relationship can be expressed using the formula *Y*=19.2262 *X* + 95.95332.

Food processing can greatly affect Se content in food, and increasing Se content in grain through foliar Se fortification is an effective way to improve Se content in wheat-derived foods.

### 3.4 Se bioavailability

Not all the Se obtained from foods can be used by the human body. The ratio of body absorbed Se to total intake Se is called bioavailability^10^. The Se from food can be effectively used by the human body only through the absorption in blood circulation or tissues and then transformed into active substances. A large difference was observed for the bioavailability of different Se types. Se in wheat-derived foods mainly can be divided into inorganic Se (i.e., selenite) and organic Se (i.e., SeMet). Organic SeMet is the main form in wheat grain, flour, and wheat-derived foods, which accounts for 70%–90%^14, 84^, 59%–82%^43, 78, 85, 86^, and 42%–83%^43, 57, 85^ of total Se, respectively.

Different forms of Se are digested and absorbed in different ways, causing different bioavailability. Organic Se is the mainly forms which can be actively absorbed by human small intestine, and the bioavailability generally fluctuates between 70% and 90%^43, 57, 85^. Inorganic Se can be absorbed via simple absorption (selenite) and passive diffusion (selenite), and the bioavailability is generally less than 50%^87^. Moreover, the long-term consumption of inorganic Se may result in toxic and side effects^10^. The molecular of H_2_Se is the key in the Se metabolism of human body. The role of H_2_Se mainly includes : 1) H_2_Se first transforms to selenite and SeMet form, and then synthesizes selenoprotein in liver and 2) SeMet and SeMeCys catalyze H_2_Se to form different Se ions forms under continuous methylation, which are discharged through respiration, urine, and sweat^88, 89^.

Se bioavailability in human body can be predicted via *in vitro* simulated gastrointestinal digestion model^88, 89^, Caco-2 based cell model^91^, animal models^92^, and human Se bioavailability detection^93^. Se bioavailability in wheat was 83% in rat feeding trials, while those in mushroom, tuna, and beef kidneys were 5%, 57%, and 97%, respectively^92^. Se bioavailability in wheat flour was also very high, which values was 75.4%–91.8%^94^. The consecutive consumption of Se-enriched wheat for six weeks could significantly increase human serum Se content, but Se-enriched fish had no obvious effect^93^. Generally, SeMet can be effectively absorbed by human body, and its bioavailability may exceed 90%. By using *in vitro* gastrointestinal digestion method to simulate bread digestion, soluble Se content after gastric and gastrointestinal digestion accounted for 75% and 89% of total Se content in bread, respectively, and SeMet accounted for 95.5% and 96.8% of soluble Se^93^.

Se bioavailability in foods is 10%–85%^10^. These differences in bioavailability might be ascribed to the following factors: 1) Se type in food - different Se types showed different bioavailability. For instance, SeMet and SeMeCys have a bioavailable of up to 90%^92^. The bioavailability of SeMet is approximately 1.5-fold higher than that of selenite^96^. 100% of selenate can be taken up, but a significant fraction will be excreted in the urine. Yet, the direct absorption of selenite is approximately 50%–60%^97^. 2) Food type - a high-fat diet can hinder Se absorption, resulting in low Se bioavailability^90^. 3) Food processing - heat treatment not only affects total Se content but also reduced Se antagonist content. However, it’s worth noting that proper heat treatment might improve Se bioavailability too^92^. 4) Antagonism with other minerals - Ca intake in the human body would reduce Se bioavailability. Meanwhile, Vc can transform selenate into insoluble Se compounds in a short time, thus reducing Se bioavailability^98^. 5) Digestion and absorption capacity of human body. For instance, gastrointestinal diseases will reduce Se bioavailability^98^.

The reports above indicated that wheat-related foods were effective Se source for people because most of the Se in wheat-related foods are organic SeMet which has high bioavailability. So, improving Se content in the wheat grain by foliar Se fortification is the key step and effective way to supplement Se in human body in most of the Se deficient areas.

### 3.5 Suitable level of Se in wheat grain and in fertilizer applied in the field

Both insufficient and excessive Se intake will have adverse effects on human health. Therefore, it should be seriously considered about the suitable Se level in the grain of wheat and the proper Se fertilizer quantity in the field which can be determined according to some important factors including the proper Se intake quantity per capita, dietary pattern, wheat Se accumulation efficiency and Se loss in food processing etc.

The range of Se intake recommended by WHO is 50–400 μg/day and this can be the reference to determine the proper Se intake quantity per capita of people. Dietary pattern, which refers to the combination of various forms of food ingredients used in people’s actual life, should also be considered^100, 101^. The representative dietary patterns are divided into Western, Oriental, and Mediterranean dietary patterns. Western dietary patterns are mainly characterized by the intake of large amounts of red meat, processed meat products, refined cereals, desserts, French fries, and high-fat dairy products. Oriental dietary patterns are mainly characterized by the intake of a large number of fruits, vegetables, legumes, whole wheat, an appropriate amount of fish, and poultry. Mediterranean dietary patterns are mainly characterized by the intake of monounsaturated fatty acids rich in olive and olive oil, whole grains, fruits, vegetables and dairy products, weekly intake of fish, poultry, nuts, and legumes. The distribution law of food Se content is as follows^62^: fish > meat > egg > cereal > fruits and vegetables. Hence, the actual Se intake per capita in western dietary patterns countries could reach the recommended level. The actual Se intake per capita in some Mediterranean dietary pattern countries also could reach the recommended level, such as Italy, but some countries such as Spain cannot not reach the recommended level. While the actual Se intake per capita in Oriental dietary patterns countries, such as China and India, did not reach the recommended level.

With wheat as Se source, the level of Se needed in wheat grains and the amount of Se to be applied in the field should be determined.

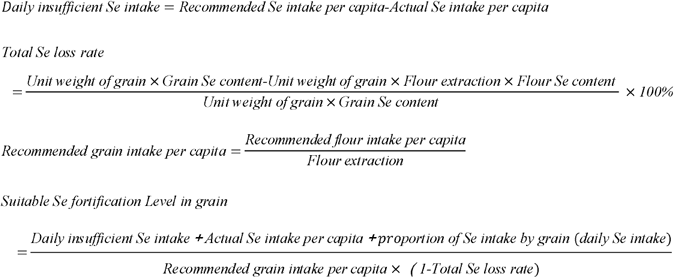

The recommended Se intake per capita is 60–400 μg·day^−1^, the recommended Se in grain is 150 g (assuming all for wheat), the actual Se intake is 43.90 μg·kg^−1^, and the grain daily Se intake accounts for approximately 22.6% in China according to the national nutrition and health guidelines. The upper limit of Se content in wheat grain is 300 μg·kg^−1^, and the scope of Se content in wheat grain is 242.74–300.00 μg·kg^−1^ in China. According to the formula and the parameters, the amount of Se fortification by foliar Se fertilizer in China should be 18.53–23.96 g·hm^−2^ to meet people’s daily need. The result varied widely between different countries, for example, the amount of Se fortification should be 2.65–3.37 and 3.93–9.88 g·hm^−2^ in India and Spain, respectively (Figure 7, Table 1 ^8, 98, 102–107^).

**Table 1.** Recommended amount of wheat Se fertilizer in different countries.

| Dietary pattern | Country | Recommended daily Se intake per capita ($\mu\text{g}\cdot\text{d}^{-1}$) | Wheat grain Se content ($\mu\text{g}\cdot\text{kg}^{-1}$) | Actual Se intake per capita ($\mu\text{g}\cdot\text{d}^{-1}$) | Recommended wheat grain Se content ($\mu\text{g}\cdot\text{kg}^{-1}$) | Recommended Se fertilizer applying amount ($\text{g}\cdot\text{hm}^{-2}$) |
| --- | --- | --- | --- | --- | --- | --- |
| Oriental dietary patterns | China | 60.00<sup>100</sup> | 47.46 | 43.90<sup>97</sup> | 242.74-300.00 | 18.53-23.96 |
|  | India | 50.00<sup>8</sup> | 171.70 | 48.00<sup>98</sup> | 272.76-300.00 | 2.65-3.37 |
| Mediterranean dietary patterns | Spain | 50.00<sup>8</sup> | 93.90 | 49.24<sup>95</sup> | 175.84-300.00 | 3.93-9.88 |
|  | Italy | 50.00<sup>8</sup> | 133.44 | 66.53<sup>96</sup> | No need to fortification | No need to fortification |
| Western dietary patterns | Australia | 55.00<sup>92</sup> | 191.83 | 57.00<sup>99</sup> | No need to fortification | No need to fortification |
|  | America | 55.00<sup>92</sup> | 304.95 | 80.00<sup>98</sup> | No need to fortification | No need to fortification |

## 4 CONCLUSION AND FUTURE TRENDS

All the reports now showed that Se nutrition enhancement of human body through the consumption of wheat in the diet is an effective and feasible way. But the effects were mainly influenced by three important processes, including 1) the acquisition of Se-rich wheat grains, 2) reduction of Se loss in the process of flour milling and food processing, 3) the improvement of bioavailability of Se in food. There were many confirmed conclusions we can make and also many further researches needed to conduct in the future based on the above three aspects.

1. The acquisition of Se-rich wheat grains. It was the most important step in Se fortification. Application of Se foliar fertilizer is the first choice which is more effective and economic to get Se-rich grain than soil fertilization according to the available reports. The application of foliar fertilizer 750 kg·hm^−2^ with Se concentration 40 mg·L^−1^ at the early filling stage can most likely make grain Se content reach 200–300 μg·kg^−1^. How to improve Se efficiency of soil in the cultivation of wheat is another aspect needed further concentration. It was necessary to further investigate the relationship between soil properties and Se bioavailability for the plants in the soil, and these research combined with other special methods might be helpful to improve Se efficiency of soil. In addition, the wheat varieties are another important factor which should be considered seriously. Many reports confirmed the large variance between different wheat varieties^33^. The grain Se content of the varieties with high Se accumulation potential was 742.33% higher than that of wheat varieties with low Se accumulation^8^. Therefore, it’s necessary to selection, breeding and application of wheat varieties with strong Se enrichment ability. Some genes/varieties related with high accumulation of Se in grain should be considered in breeding^8^. Few investigations were conducted on the effects of other nutrients on wheat Se accumulation such as foliar Se fortification increased grain Se and Zn content, but reduced grain Cu, Fe, K, Mg and S content^109^.
2. Reduction of Se loss in the process of flour milling and food processing. Flour Se content can be retained to the maximum extent when the peeling rate is 4%–5% and the milling ratio is 70%–80%, and try to choose low temperature method to keep Se content in wheat derived-foods according to the available reports. But few reports focused on the mechanism of the effects on Se loss in different food processing methods.
3. The improvement of bioavailability of Se in food. The metabolism and regulation of different forms of Se in body and the interaction mechanism with other substances needed further investigation. The information will contribute to the other ways to supplement Se of human body. For example, the addition of food additives can improve Se bioavailability in food. This process should also be considered in the food industry.

Se deficiency is a worldwide problem. It is an effective and feasible way to supplement Se for people through Se fortification in wheat. We can adopt the strategy including the above three aspects to get enough Se from wheat. But this strategy needs more public attention, more research and application results to achieve this goal.

## 5 CONFILCT OF INTEREST

The authors declare that the research was conducted in the absence of any commercial or financial relationships that could be construed as a potential conflict of interest.

## 6 AUTHOR CONTRIBUTIONS

MW, SL and ZWS collected and analysed data. MW, FMK and XCZ wrote this paper. XCZ, FMK and BQL conceived and modified this paper.

## 7 ACKNOWLEDGMENTS

This paper was supported by Agricultural Variety Improvement Project of Shandong Province (2019LZGC001) and Special Project of Regional Science and Technology Collaborative Innovation between District Science and Technology Department and Rikaze Municipal Government in 2020 (Research and Demonstration Application of Bioaugmentation Technology for trace nutrients of highland barley in Rikaze area).

