## Supplemental Tables for "Selenium in wheat from farming to food"

**Table S1** Selenium content in wheat grains around the world

| **Serial number** | **Country** | **Fertilizing method** | **Fertilizing amount** | **Number of samples** | **Grain Selenium content (μg·kg^-1^)** | | **References** |
| --- | --- | --- | --- | --- | --- | --- | --- |
|  |  |  |  |  | **Mean** | **Range** |  |
| 1 | Australia | ND | ND | 7 | 442.86 | 100.00–700.00 | Robinson, 1936 ^50^ |
|  | Argentina | ND | ND | 4 | 600.00 | 400.00–800.00 | Robinson, 1936 ^50^ |
|  | Canada | ND | ND | 1 | 1900.00 | ND | Robinson, 1936 ^50^ |
|  | Mexico | ND | ND | 1 | 600.00 | ND | Robinson, 1936 ^50^ |
|  | South Africa | ND | ND | 2 | 850.00 | 200.00–1500.00 | Robinson, 1936 ^50^ |
|  | Spain | ND | ND | 4 | 500.00 | 200.00–800.00 | Robinson, 1936 ^50^ |
|  | America | ND | ND | 6 | 183.33 | 100.00–300.00 | Robinson, 1936 ^50^ |
|  | New Zealand | ND | ND | 1 | 400.00 | ND | Robinson, 1936 ^50^ |
|  | Hungary | ND | ND | 2 | 350.00 | 300.00–400.00 | Robinson, 1936 ^50^ |
| 2 | America | ND | ND | 190 | 401.80 | 144.00–789.00 | Harn et al.,1981 ^49^ |
| 3 | Norway | Soil fertilization | 5.82/6.45/12.9 g Se·hm^-2^ | 47 | 249.19 | 25.00–497.00 | Tveitnes et al.,1995 ^48^ |
|  | Norway | ND | ND | 16 | 26.75 | 5.00–94.00 | Tveitnes et al.,1995 ^48^ |
| 4 | Spain | ND | ND | 2 | 35.50 | 31.70–39.40 | Diaz-Alarcon et al., 1996 ^46^ |
| 5 | Mexico | ND | ND | 9 | 190 | 140.00–210.00 | Wyatt et al., 1996 ^47^ |
| 6 | Australia | ND | ND | 2 | 460 | 410.00–510.00 | Lyons et al., 2005 ^45^ |
|  | Australia | ND | ND | 151 | 147.59 | 15.00–520.00 | Lyons et al., 2005 ^45^ |
| 7 | Italy | ND | ND | 749 | 123.5 | 7.00–245.00 | Spadoni et al., 2007 ^44^ |
| 8 | India | ND | ND | 17 | 165.76 | 107.00–272.00 | Yadav et al., 2008 ^43^ |
| 9 | Italy | ND | ND | 3 | 109.17 | 95.20–122.50 | Cubadda et al., 2009 ^41^ |
| 10 | Slovakia | ND | ND | 4 | 46.25 | 30.00–60.00 | Ducsay et al., 2009 ^42^ |
|  | Slovakia | Soil fertilization | 150/300/500 g Se·hm^-2^ | 6 | 526.83 | 140.00–818.00 | Ducsay et al., 2009 ^42^ |
| 11 | China | ND | ND | 1 | 38.10 | ND | Tang et al.,2010 ^39^ |
|  | China | Soaking seeds | 0.1/0.5/1.0/2.5 mg Se·L^-1^ | 1 | 70.50 | 64.80–77.60 | Tang et al.,2010 ^39^ |
| 12 | China | ND | ND | 62 | 35.47 | 16.00–72.00 | Lu et al.,2010 ^40^ |
| 13 | China | ND | ND | 1 | 78.34 | 90.00–100.00 | Tang et al., 2011 ^38^ |
|  | China | Soil fertilization | 0.1/0.25/0.5/1.0/2.0/3.0/5.0 mg·kg^-1^ Na_2_SeO_3_.5H_2_O | 8 | 2447.50 | 210.00–6090.00 | Tang et al., 2011 ^38^ |
|  | China | Foliar fertilization | 15/30/45/60/75 g Se·hm^-2^ | 10 | 37.84 | 180.00–1060.00 | Tang et al., 2011 ^38^ |
| 14 | China | Foliar fertilization | 57.95 g Se·hm^-2^ | 1 | 288.94 | ND | Zhang et al., 2012 ^37^ |
|  | China | ND | ND | 3 | 114.16 | 102.16–121.49 | Zhang et al., 2012 ^37^ |

**Continued Table S1** Selenium content in wheat grains around the world

| **Serial number** | **Country** | **Fertilizing method** | **Fertilizing amount** | **Number of samples** | **Grain Selenium content (μg·kg^-1^)** | | **References** |
| --- | --- | --- | --- | --- | --- | --- | --- |
|  |  |  |  |  | **Mean** | **Range** |  |
| 15 | Portugal | Foliar fertilization | 4/20/100 g Se·hm^-2^ | 12 | 466.67 | 75.00–2100.000 | Galinha et al., 2012 ^52^ |
| 16 | China | ND | ND | 1 | 12.50 | 8.60–17.10 | Zhang et al., 2012 ^36^ |
|  | China | Foliar fertilization | 57.95 g Se·hm^-2^ | 1 | 755.90 | ND | Zhang et al., 2012 ^36^ |
| 17 | China | ND | ND | 4 | 97.50 | 71.00–131.00 | Zhang et al., 2012 ^35^ |
|  | China | Foliar fertilization | 3.626 g Se·hm^-2^ | 2 | 141.50 | 129.00–154.00 | Zhang et al., 2012 ^35^ |
| 18 | China | ND | ND | 3 | 7.67 | 5.00–12.00 | Wang，2012 ^34^ |
|  | China | Foliar fertilization | 3/15/19/30/38/46/57/61/76 g Se·hm^-2^ | 8 | 484.13 | 38.00–1006.00 | Wang，2012 ^34^ |
|  | China | Soil fertilization | 150/300/450/600 g Se·hm^-2^ | 4 | 166.25 | 84.00–339.00 | Wang，2012 ^34^ |
| 19 | Algeria | ND | ND | 8 | 52.00 | 21.99–153.00 | Beladel et al., 2013 ^33^ |
| 20 | China | ND | ND | 1 | 177.34 | ND | Pei et al., 2013 ^33^ |
| 21 | Brazil | Soaking seeds | 10 μM Na_2_SeO_4_ | 20 | 245.00 | 190.00–300.00 | Souza et al., 2014 ^55^ |
| 22 | China | Foliar fertilization | 57.95 g Se·hm^-2^ | 2 | 330.07 | 192.69–467.45 | Meng，2014 ^31^ |
|  | China | ND | ND | 6 | 58.15 | 30.63–96.76 | Meng，2014 ^31^ |
| 23 | Spain | ND | ND | 1 | 66.60 | ND | Poblaciones et al., 2014 ^30^ |
|  | Spain | Foliar fertilization | 10/20/40 g Se·hm^-2^ | 1 | 551.47 | 153.60–1383.20 | Poblaciones et al., 2014 ^30^ |
| 24 | China | ND | ND | 3 | 101.33 | 31.00–144.00 | Zhang et al., 2015 ^27^ |
|  | China | Foliar fertilization | 0.0089 g Se·hm^-2^ | 3 | 1248.00 | 803.00–1694.00 | Zhang et al., 2015 ^27^ |
| 25 | China | ND | ND | 3 | 29.00 | 19.00–38.00 | Yu,2015 ^28^ |
|  | China | Foliar fertilization | 18/45 g Se·hm^-2^ | 6 | 139.17 | 26.00–397.00 | Yu,2015 ^28^ |
|  | China | Soil fertilization | 15/700 g Se·hm^-2^ | 6 | 166.83 | 23.00–256.00 | Yu,2015 ^28^ |
| 26 | Portugal | ND | ND | 9 | 85.00 | 60.00–150.00 | Galinha et al., 2015 ^29^ |
|  | Portugal | Foliar fertilization | 100 g Se·hm^-2^ | 7 | 1331.67 | 610.00–2700.00 | Galinha et al., 2015 ^29^ |
|  | Portugal | Soil fertilization | 100 g Se·hm^-2^ | 7 | 1111.33 | 760.00–1510.00 | Galinha et al., 2015 ^29^ |
| 27 | China | Foliar fertilization | 120 g Se·hm^-2^ | 1 | 1640.00 | ND | Wang et al., 2016 ^13^ |
| 28 | China | ND | ND | 31 | 31.00 | 0.00–117.00 | Liu et al., 2016 ^8^ |
|  | China | Foliar fertilization | 57.95 g Se·hm^-2^ | 31 | 647.80 | 152.6–1796.5 | Liu et al., 2016 ^8^ |
| 29 | Italy | ND | ND | 5 | 148.75 | 100.00–220.00 | De Vita et al., 2017 ^26^ |
|  | Italy | Foliar fertilization | 1/5/10/15/20/25/50/80/100/120 g Se·hm^-2^ | 4 | 3864.00 | 330.00–8270.00 | De Vita et al., 2017 ^26^ |
| 30 | China | Foliar fertilization | 30/60 g Se·hm^-2^ | 1 | 225.00 | 70.00–380.00 | Wang et al.,2017 ^57^ |
| 31 | China | ND | ND | 7 | 52.57 | 27.00–143.00 | Wang et al., 2018 ^23^ |

**Continued Table S1** Selenium content in wheat grains around the world

| **Serial number** | **Country** | **Fertilizing method** | **Fertilizing amount** | **Number of samples** | **Grain Selenium content (μg·kg^-1^)** | | **References** |
| --- | --- | --- | --- | --- | --- | --- | --- |
|  |  |  |  |  | **Mean** | **Range** |  |
| 31 | China | Foliar fertilization | 22.49 g Se·hm^-2^ | 7 | 315.86 | 58.00–608.00 | Wang et al., 2018 ^23^ |
| 32 | Israel | ND | ND | 1 | 128.22 | ND | Ji et al., 2018 ^22^ |
|  | Russia | ND | ND | 1 | 125.48 | ND | Ji et al., 2018 ^22^ |
|  | Canada | ND | ND | 13 | 97.81 | 30.86–144.77 | Ji et al., 2018 ^22^ |
|  | Portugal | ND | ND | 1 | 120.37 | ND | Ji et al., 2018 ^22^ |
|  | Australia | ND | ND | 1 | 141.27 | ND | Ji et al., 2018 ^22^ |
|  | Ukraine | ND | ND | 1 | 164.02 | ND | Ji et al., 2018 ^22^ |
|  | South Africa | ND | ND | 1 | 148.34 | ND | Ji et al., 2018 ^22^ |
|  | America | ND | ND | 3 | 115.61 | 100.98–123.45 | Ji et al., 2018 ^22^ |
|  | Argentina | ND | ND | 1 | 121.12 | ND | Ji et al., 2018 ^22^ |
|  | Spain | ND | ND | 2 | 82.39 | 58.36–106.41 | Ji et al., 2018 ^22^ |
|  | Cypress | ND | ND | 1 | 55.06 | ND | Ji et al., 2018 ^22^ |
|  | Italy | ND | ND | 1 | 30.86 | ND | Ji et al., 2018 ^22^ |
|  | Syria | ND | ND | 1 | 144.77 | ND | Ji et al., 2018 ^22^ |
|  | Iraq | ND | ND | 1 | 97.17 | ND | Ji et al., 2018 ^22^ |
|  | India | ND | ND | 1 | 61.06 | ND | Ji et al., 2018 ^22^ |
| 33 | China | ND | ND | 3 | 47.99 | 44.04–50.75 | Feng et al., 2018 ^24^ |
|  | China | Foliar fertilization | 15/30/45/60 g Se·hm^-2^ | 12 | 288.80 | 177.19–428.21 | Feng et al., 2018 ^24^ |
| 34 | China | ND | ND | 6 | 53.83 | 26.00–72.00 | Jin et al., 2018 ^25^ |
|  | China | Foliar fertilization | 2.5/5/7.5/10/12.5 g Se·hm^-2^ | 24 | 199.42 | 87.00–443.00 | Jin et al., 2018 ^25^ |
| 35 | China | Foliar fertilization | 57.95 g Se·hm^-2^ | 4 | 2081.10 | 1810.40–2515.50 | Zhang et al., 2019 ^16^ |
|  | China | ND | ND | 4 | 67.75 | 58.00–75.00 | Zhang et al., 2019 ^16^ |
| 36 | China | ND | ND | 5 | 307.20 | 214.00–396.00 | Qin et al., 2019 ^17^ |
| 37 | China | ND | ND | 1 | 28.56 | ND | Dong et al., 2019 ^18^ |
| 38 | China | ND | ND | 1 | 225.88 | ND | Chen, 2019 ^19^ |
|  | China | Soil fertilization | 425 g Se·hm^-2^ | 1 | 375.04 | ND | Chen, 2019 ^19^ |
|  | China | Foliar fertilization | 45 g Se·hm^-2^ | 1 | 748.47 | 736.21–1621.73 | Chen, 2019 ^19^ |
| 39 | China | ND | ND | 2 | 39.00 | 26.00–45.00 | Zou et al., 2019 ^20^ |
|  | China | Foliar fertilization | 1 g Se·hm^-2^ | 2 | 197.00 | 144.00–297.00 | Zou et al., 2019 ^20^ |
|  | India | ND | ND | 2 | 188.17 | 26.00–549.00 | Zou et al., 2019 ^20^ |

**Continued Table S1** Selenium content in wheat grains around the world

| **Serial number** | **Country** | **Fertilizing method** | **Fertilizing amount** | **Number of samples** | **Grain Selenium content (μg·kg^-1^)** | | **References** |
| --- | --- | --- | --- | --- | --- | --- | --- |
|  |  |  |  |  | **Mean** | **Range** |  |
| 39 | India | Foliar fertilization | 1 g Se·hm^-2^ | 2 | 406.17 | 178.00–725.00 | Zou et al., 2019 ^20^ |
|  | Mexico | ND | ND | 2 | 24.50 | 16.00–33.00 | Zou et al., 2019 ^20^ |
|  | Mexico | Foliar fertilization | 1 g Se·hm^-2^ | 2 | 199.00 | 185.00–213.00 | Zou et al., 2019 ^20^ |
|  | Turkey | ND | ND | 1 | 41.00 | 4.00–66.00 | Zou et al., 2019 ^20^ |
|  | Turkey | Foliar fertilization | 1 g Se·hm^-2^ | 1 | 238.00 | 155.00–244.00 | Zou et al., 2019 ^20^ |
|  | Pakistan | ND | ND | 1 | 125.20 | 31.00–257.00 | Zou et al., 2019 ^20^ |
|  | Pakistan | Foliar fertilization | 1 g Se·hm^-2^ | 1 | 302.80 | 222.00–446.00 | Zou et al., 2019 ^20^ |
|  | South Africa | ND | ND | 2 | 55.75 | 20.00–86.00 | Zou et al., 2019 ^20^ |
|  | South Africa | Foliar fertilization | 1 g Se·hm^-2^ | 2 | 643.00 | 558.0–711.00 | Zou et al., 2019 ^20^ |
| 40 | Spain | ND | ND | ND | 27.96 | 14.68–51.84 | López-Bellido et al., 2019 ^21^ |
| 41 | China | Foliar fertilization | 20/100 g Se·hm^-2^ | 4 | 3442.50 | 610.00–68200.00 | Wang et al., 2020 ^15^ |
|  | China | ND | ND | ND | 120.00 | ND | Wang et al., 2020 ^15^ |
| - | Mean | - | - | - | 411.38 | 0.00**-**68200.00 | - |

Note: ND indicates that relevant information is not mentioned in the literature.

**Table S2** Selenium content in different components of wheat

| **Serial number** | **Country** | **Peeling rate (%)** | **Flour extraction rate (%)** | **Number of samples** | **Selenium content in wheat flour (μg·kg^-1^)** | | **Selenium content in wheat Brans (μg·kg^-1^)** | | **References** |
| --- | --- | --- | --- | --- | --- | --- | --- | --- | --- |
|  |  |  |  |  | **Mean** | **Range** | **Mean** | **Range** |  |
| 1 | Canada | ND | ND | 3 | 280.00 | 260.00–280.00 | ND | ND | Arthur, 1972 ^66^ |
|  | Canada | ND | 100% | 3 | 610.00 | 560.00–650.00 | ND | ND | Arthur, 1972 ^66^ |
| 2 | Germany | ND | ND | 2 | 24.00 | 23.00–25.00 | ND | ND | Oster et al., 1989 ^65^ |
|  | New Zealand | ND | ND | 2 | 18.00 | 14.00–22.00 | ND | ND | Oster et al., 1989 ^65^ |
|  | Greece | ND | ND | 2 | 189.00 | 38.00–340.00 | ND | ND | Oster et al., 1989 ^65^ |
|  | America | ND | ND | 1 | 190.00 | ND | ND | ND | Oster et al., 1989 ^65^ |
| 3 | Britain | ND | 100% | 23 | 59.00 | 23.00–110.00 | ND | ND | Barclay et al., 1995 ^64^ |
|  | Britain | ND | ND | 47 | 38.00 | 17.00–85.00 | ND | ND | Barclay et al., 1995 ^64^ |
| 4 | Spain | ND | ND | 1 | 32.20 | ND | ND | ND | Diaz-Alarcon et al., 1996 ^63^ |
| 5 | Slovakia | ND | ND | 11 | 25.10 | 15.00–32.30 | ND | ND | Kadrabova et al., 1997 ^67^ |
| 6 | Ireland | ND | ND | ND | 64.00 | 13.00–99.00 | ND | ND | Murphy, 2001 ^62^ |
|  | Ireland | ND | 100% | 3 | 130.00 | 128.00–132.00 | ND | ND | Murphy, 2001 ^62^ |
| 7 | Croatia | ND | ND | 2 | 57.55 | 56.30–58.80 | ND | ND | Klapec et al., 2004 ^61^ |
| 8 | Australia | ND | 60% | 2 | 170.50 | 41.00–300.00 | 235.00 | 60.00–410.00 | Lyons et al., 2005 ^45^ |
| 9 | Libya | ND | ND | 1 | 45.00 | ND | ND | ND | Alamin et al., 2006 ^60^ |
| 10 | China | ND | ND | 3 | 67.33 | 14.00–120.00 | ND | ND | Wang et al., 2008 ^59^ |
| 11 | Italy | ND | 66% | 3 | 91.27 | 83.50–97.30 | ND | ND | Cubadda et al., 2009 ^41^ |
| 12 | Spain | ND | ND | 3 | 55.67 | 50.00–67.00 | ND | ND | Matos-Reyes et al., 2010 ^58^ |
| 13 | Britain | ND | ND | 2 | 553.46 | 30.30–2008.20 | ND | ND | Hart et al., 2011 ^57^ |
|  | Britain | ND | 100% | 2 | 640.31 | 41.60–2259.10 | ND | ND | Hart et al., 2011 ^57^ |
| 14 | Spain | ND | ND | 1 | 391.60 | 44.00–1235.60 | ND | ND | Poblaciones et al., 2014 ^30^ |
| 15 | China | ND | 100% | 6 | 102.35 | 64.30–118.00 | ND | ND | Gao et al., 2015 ^68^ |
| 16 | China | ND | 89% | 15 | 68.27 | 19.00–288.00 | 119.93 | 15.00–520.00 | Yu,2015 ^28^ |
| 17 | China | ND | 73.22% | 1 | 1420.00 | ND | 1910 | ND | Wang et al., 2016 ^13^ |
| 18 | China | 0-15.2% | ND | 2 | 204.21 | 20.00–580.00 | ND | ND | Li et al., 2017 ^10^ |
|  | China | 0-14.8% | ND | 1 | 411.00 | 110.00–850.00 | ND | ND | Li et al., 2017 ^10^ |
| 19 | Italy | ND | 100% | 1 | 2640.00 | 150.00–5380.00 | ND | ND | De Vita et al., 2017 ^26^ |
|  | Italy | ND | 66% | 1 | 2343.33 | 120.00–4690.00 | 3113.33 | 230.00–6360.00 | De Vita et al., 2017 ^26^ |
| 20 | China | ND | ND | 1 | 795.96 | 242.01–1433.81 | 932.03 | 236.61–1715.52 | Chen, 2019 ^19^ |
| - | Mean | - | - | - | 404.04 | 13.00–5380.00 | 1547.59 | 60–6360 | **-** |

Note: ND indicates that relevant information is not mentioned in the literature.

**Table S3** Selenium content in different wheat processed foods

| **Name** | **Number of samples** | **Selenium content (μg·kg^-1^)** | | **References** | **Remark** |
| --- | --- | --- | --- | --- | --- |
|  |  | **Mean** | **Range** |  |  |
| Bread | 3 | 676.67 | 650.00–710.00 | Arthur, 1972 ^66^ | Whole wheat |
|  | 3 | 536.67 | 420.00–670.00 | Arthur, 1972 ^66^ | White bread |
|  | 8 | 19.00 | 13.00–25.00 | Oster et al., 1989 ^65^ | White bread |
|  | 16 | 220 | 140.00–270.00 | Eurola et al., 1991 ^78^ |  |
|  | 1 | 203.00 | ND | Aro et al., 1994 ^77^ |  |
|  | 5 | 180.20 | 87.00–337.00 | Aro et al., 1994 ^77^ |  |
|  | 25 | 92.00 | 26.00–200.00 | Barclay et al., 1995 ^64^ | Whole wheat |
|  | 24 | 44.00 | 18.00–35.00 | Barclay et al., 1995 ^64^ |  |
|  | 5 | 42.12 | 25.50–77.90 | Diaz-Alarcon et al., 1996 ^63^ |  |
|  | 3 | 23.63 | 19.50–26.70 | Finley et al., 1996 ^76^ |  |
|  | 3 | 27.97 | 22.70–32.60 | Finley et al., 1996 ^76^ | Whole wheat |
|  | 6 | 17.60 | 14.30–21.50 | Kadrabova et al.,1997 ^67^ |  |
|  | 4 | 127.10 | 63.99–274.00 | Hussein et al., 1999 ^81^ |  |
|  | 1 | 364.29 | ND | Holben et al., 1999 ^75^ | Whole wheat |
|  | 1 | 284.00 | ND | Holben et al., 1999 ^75^ | White bread |
|  | 26 | 129.00 | 84.00–158.00 | Murphyt et al., 2001 ^63^ | Whole wheat |
|  | 18 | 66.00 | 55.00–95.00 | Murphyt et al., 2001 ^62^ | White bread |
|  | 20 | 118.00 | 94.00–146.00 | Murphyt et al., 2001 ^62^ | Adding bran |
|  | 2 | 57.55 | 45.70–47.70 | Klapec et al., 2004 ^61^ |  |
|  | 1 | 171.33 | 167.00–177.00 | Lyons et al., 2005 ^45^ | White bread |
|  | 1 | 172.33 | 170.00–177.00 | Lyons et al., 2005 ^45^ | Whole wheat |
|  | ND | 91.90 | 70.00–131.8 | Pappa et al., 2006 ^74^ |  |
|  | 1 | 40 | ND | Bryszewska et al., 2007 ^73^ |  |
|  | 16 | 578.99 | 34.40–2090.50 | Hart et al., 2011 ^57^ | Soil fertilization |
|  | 16 | 669.44 | 39.60–2372.20 | Hart et al., 2011 ^57^ | Whole wheat+Soil fertilization |
| Pasta | 6 | 821.67 | 650.00–1420.00 | Arthur, 1972 ^66^ |  |
|  | 7 | 390.00 | ND | Olson et al., 1984 ^79^ |  |
|  | 1 | 870.00 | ND | Olson et al., 1984 ^79^ | Whole wheat |
|  | 10 | 55.00 | 35.00–88.00 | Barclay et al., 1995 ^64^ | Whole wheat |
|  | 12 | 48.00 | 37.00–58.00 | Barclay et al., 1995 ^64^ |  |
|  | 2 | 24.40 | 22.30–26.80 | Diaz-Alarcon et al., 1996 ^63^ |  |

**Continued Table S3** Selenium content in different wheat processed foods

| **Name** | **Number of samples** | **Selenium content (μg·kg^-1^)** | | **References** | **Remark** |
| --- | --- | --- | --- | --- | --- |
|  |  | **Mean** | **Range** |  |  |
| Pasta | 1 | 32.70 | ND | Finley et al., 1996 ^76^ |  |
|  | 1 | 212.86 | ND | Holben et al., 1999 ^75^ |  |
|  | 20 | 66.00 | 49.00–78.00 | Murphyt et al., 2001 ^62^ | Fresh + whole wheat |
|  | 6 | 221.17 | 48.00–526.00 | Alamin et al., 2006 ^60^ |  |
|  | 3 | 71.00 | 20.00–130.00 | Wang et al., 2008 ^59^ |  |
|  | 3 | 91.10 | 83.50–97.30 | Cubadda et al., 2009 ^41^ |  |
|  | 3 | 88.37 | 80.30–92.80 | Cubadda et al., 2009 ^41^ |  |
|  | 12 | 154.00 | 84.00–227.00 | Cubadda et al., 2009 ^41^ |  |
|  | 7 | 367.00 | 22.40–1190.40 | Poblaciones et al., 2014 ^30^ |  |
|  | 1 | 23.92 | ND | Wang et al., 2015 ^72^ |  |
|  | 1 | 492.10 | ND | Wang et al., 2015 ^72^ | Adding selenium-rich yeast |
| Steamed buns | 1 | 770.00 | ND | Wang et al., 2016 ^13^ | Foliar fertilization |
| Noodle | 3 | 936.67 | 760.00–1130.00 | Arthur, 1972 ^66^ |  |
|  | 2 | 430.00 | ND | Olson et al., 1984 ^79^ | Whole wheat |
|  | 5 | 590.00 | ND | Olson et al., 1984 ^79^ | Adding eggs |
|  | 1 | 48 | ND | Oster et al., 1989 ^65^ |  |
|  | 2 | 29.50 | 27.00–32.10 | Diaz-Alarcon et al., 1996 ^63^ |  |
|  | 1 | 23.40 | ND | Finley et al., 1996 ^76^ |  |
|  | 3 | 56.80 | 53.00–59.00 | Kadrabova et al., 1997 ^67^ |  |
|  | 1 | 216.88 | ND | Holben et al., 1999 ^75^ | Adding eggs |
|  | 4 | 43.80 | ND | Klapec et al., 2004 ^61^ |  |
|  | 4 | 118.25 | 15.00–220.00 | Wang et al., 2008 ^59^ |  |
| Wheat groats | 3 | 40.00 | 30.00–50.00 | Arthur, 1972 ^66^ |  |
|  | 2 | 46 | 43.00–49.00 | Ferretti et al, 1974 ^80^ |  |
|  | 5 | 51.00 | ND | Olson et al., 1984 ^79^ |  |
|  | 28 | 31.00 | 28.00–34.00 | Murphyt et al., 2001 ^62^ |  |
| Biscuits | 14 | 60.00 | ND | Olson et al., 1984 ^79^ |  |
|  | 15 | 242.00 | 90.00–600.00 | Arthur, 1972 ^66^ |  |
|  | 4 | 60 | 50.00–60.00 | Eurola et al., 1991 ^78^ |  |
|  | 3 | 28.20 | 8.20–44.90 | Diaz-Alarcon et al., 1996 ^63^ |  |
|  | 2 | 4.50 | 3.60–5.40 | Finley et al., 1996 ^76^ |  |

**Continued Table S3** Selenium content in different wheat processed foods

| **Name** | **Number of samples** | **Selenium content (μg·kg^-1^)** | | **References** | **Remark** |
| --- | --- | --- | --- | --- | --- |
|  |  | **Mean** | **Range** |  |  |
| Biscuits | 2 | 34.20 | ND | Klapec et al., 2004 ^61^ |  |
|  | 1 | 420.00 | ND | Zhai et al., 2018 ^71^ |  |
|  | 1 | 500.00 | ND | He, 2019 ^70^ |  |
| - | Mean | 211.86 | 15.00–2372.20 | *-* | - |

Note: ND indicates that relevant information is not mentioned in the literature.
